# Scarification of Exotic and Indigenous Plant Seeds in Nigeria: Effect on Dormancy and Germination

**DOI:** 10.1101/354993

**Authors:** Amodu Emmanuel, Osuolale Olayinka

## Abstract

Dormancy is exhibited in many seed producing plants. It could be endogenous or exogenous, depending on the plant and the type of seed the plant produce. A survival strategy, plant use to conserve their genetic materials during unfavourable conditions. Scarification treatments has been used in this work to break the dormancy of *Anacardium occidentale, Annona muricata, Jatropha curcas, Tamarindus indica* and *Artocarpus heterophyllus* using 65% Nitric acid (HNO_3_), 65% Sulphuric acid (H_2_SO_4_), 0.5% Potassium tetraoxosulphate(VI) (K_2_SO_4_), 0.5% Urea (CH_4_N_2_O), 43% Ethanol (C_2_H_6_O) and Distilled water. Nitric acid (65% HNO_3_) produced the best result for *Anacardium occidentale* with high numbers of seedlings and a germination period of 15 days. *Jatropha curcas* did not produce a favourable result from the treatments. *Tamarindus indica,* water treatment produced the best result with six days of germination shorter than the controlled value (16 days). Nitric acid (65% HNO_3_) and water favor *Annona muricata* with germination period of 19 days as against 24 days for control experiment. Water and Potassium sulphate are the best treatments for *Artocarpus heterophyllus* as they produce viable seedlings with short germination period of 14 and 15 days which give a good result better than the 18 days of the control experiment.

## INTRODUCTION

Dormancy indicates a temporal retardation in growth of an organism. Dormant organism’s shows periods of arrested growth, this is highly pronounced in some animals and plants. In plants, dormancy can be observed in seed and seedling germination. According to Vleeshouwers *et al.* (1995), Thompson (2000), Fenner and Thompson (2005), dormancy is part of the characteristic requirement for the growth of plant seeds and its survival. Dormancy is exhibited by many plants as a survival strategy, this enable them to survive in harsh or extreme climates. The term dormancy is used to describe a seed that fails to germinate under favourable condition at a specified time. Seed germination process in some plants are temporarily delayed due to dormancy, this serves as an advantage for seed dispersal due to seasonal changes in different parts of the world (Hilhorst, 1995; Baskin & Baskin, 1998). Accessing the part of the seed from which dormancy can be classified, exogenous and endogenous dormancy are the two types of dormancy recognized. Exogenous dormancy is imposed by the seed coat on the matured embryo, it is caused when the seed coat is too hard, limiting the free expansion of the embryo for the protrusion of the plume and radicle during germination. The seed coat becomes impermeable during seed maturation and drying. Impermeable seed coats and other layers prevent seeds from water up take and gaseous absorption consequently preventing seed germination (Li & Foley, 1997). High concentration of growth inhibitors such as abscisic acid (C_15_H_20_O_4_) on the seed coat and its surrounding layers can delay the growth of the embryo. Endogenous dormancy is also known as embryo dormancy, this is due to some conditions embedded in the embryo such as the presence of growth inhibitors and absence of growth promoters. This will prevent embryo growth and seed germination until chemical change takes place. Seed development of dormancy, desiccation tolerance and accumulation of food reserves occur when seed have undergo embryogenesis and seed maturation phase also known as the growth phase (Bentsink & Koornneef, 2008).

Seed dormancy may be primary or secondary. Seeds exhibiting primary state of dormancy are released from their parent plant in a dormant state. Examples of families with such seeds are Anacardiaceae, Sapindaceae and Cannaceae. Seeds with secondary dormancy are released in non-dormant state but become dormant if the condition for germination becomes unfavorable. There are various causes of dormancy some of which are: Hard seed coat, Immature embryo, Germination inhibitors, Period after ripening: Some seeds have a period of ripening. Those seeds germinate only after the completion of that period. Dormancy mechanism has been diversified due to diversity of habitat and the climates of the habitat which those seeds operate. Naturally, the seed coats of physically dormant seeds becomes permeable through repeated heating and cooling over long period of time in the soil seed bank (Finch-Savage & Leubner-Metzger, 2006; Offord *et al.*, 2009). For example, temperature fluctuation during dry season in northern Australia facilitate dormancy breaking in impermeable seeds of *Stylosanthes humilis* and *Stylosanthes hamata* (Baskin, 2003).

Nikolaeva (1969) devised a dormancy classification system stressing that dormancy is determined by both morphological and physiological properties of the seed (Nikolaeva, 2004). light and gibberellins can both break dormancy and promote germination in seeds with coat dormancy according to Casal and Sánchez (1998), Sanchez and Mella (2004), and Kucera *et al.* (2005).

The abundant form of dormancy is physiological dormancy which is found in seeds of gymnosperms and most angiosperms. It is the most prevalent form of dormancy in temperate seed banks and abundant dormancy class in the field. There are three level of physiological dormancy; deep, intermediate and non-deep (Baskin & Baskin, 2004). Embryos that are differentiated into cotyledons and hypocotyl-radicle undergo morphological dormancy when the seeds are under developed in terms of embryos size. These embryos needs time to grow and germinate e.g. *Apium graveolens* (Jacobsen & Pressman, 1979). Morphophysiological dormancy common in seeds with underdeveloped embryos, but in addition they have a physiological component to their dormancy (Baskin & Baskin, 2004). These seeds therefore require a combination of warm or cold stratification and other forms of dormancy-breaking treatments which can be replaced by gibberellic acid application. Examples of plants and families with morphophysiological dormancy are *Trollius ledebouri, Fraxinus excelsior, Ranunculaceae, Oleaceae* (Finch-Savage & Clay, 1997). Mechanical or chemical scarification can break Physical dormancy caused by water-impermeable layers of palisade cells in the seed coat (Baskin, 2003).

It has been discovered that after most artificial treatments under natural condition, the first site of water penetration in seeds is the strophiole (Baskin & Baskin, 1998). Several techniques have been used to soften seed coats and make seeds permeable, some of which are: Acid scarification (soaking seeds in con. Sulphuric or Nitric acid). Which is very effective in treating Acacia seeds. Physical dormancy has been identified in the seeds of these plants across 15 angiosperm families (Rooser, 2017).

Water temperature of about 40 °C or less has been proven to be effective in promoting germination when seeds are soaked in it, but majorly in seeds with permeable seed coat. However, boiling some seed species in water removes cuticle and sometimes part of the palisade layers of the seed coat which effectively break dormancy. Soaking in water within the range 60–90 °C is often as effective as soaking at 100 °C. lower temperatures creates less chances of damage (Baskin & Baskin, 1998). According to Bewley and Black (1994), Secondary dormancy occurs in some non-dormant and post dormant seeds that are exposed to unfavorable condition for germination. Though the mechanisms of secondary dormancy are not fully understood yet, but it might involve the loss of sensitivity in receptors of the plasma membrane. Our study therefore sets out to determine the best chemical treatment in breaking seed dormancy, determine the germination period of some seeds and evaluate the effect of some chemical treatments on seed dormancy and germination.

## MATERIALS AND METHODS

The materials for this study are nursery bags, 65% Nitric acid (HNO_3_), 65% Sulphuric acid (H_2_SO_4_), 0.5% Potassium tetraoxosulphate VI (K_2_SO_4_), 0.5% Urea (CH_4_N_2_O), 43% Ethanol (C_2_H_6_O), Distilled water (H_2_O), 110 seeds of *Anacardium occidentales* Linn, 110 seeds of *Annona muricata* Linn, 110 seeds of *Jatropha curcas* Linn, 110 seeds of *Tamarindus indica* Linn, 110 seeds of *Artocarpus heterophyllus* Lam, specimen bottles, loamy soil, hand gloves and hoe.

## METHODS

The prepared chemicals were collected from the laboratory. 100 seeds each of *Anacardium occidentale* Linn, *Annona muricata* Linn, *Jatropha curcas* Linn, *Tamarindus indica* Linn, *Artocarpus heterophyllus* Lam were soaked in various chemicals (HNO_3,_ H_2_SO_4,_ H_2_O, K_2_SO_4,_ CH_4_N_2_O and C_2_H_6_O) as shown in plate 1 and 2. The seeds in HNO_3_ and H_2_SO_4_ acid were removed at an interval of 10 minutes (i.e. 10, 20, and 30 minutes respectively for the various seeds) according to the method used by El-Siddig *et al.* (2001). This method was modified by rinsing and soaking the seeds in distilled water for 24 hours. The various, seeds in equal numbers were soaked in 0.5% urea, 43% ethanol, 0.5% K_2_SO_4_, and distilled water separately for 24 hours (Yücel, 2000; Purohit *et al.*, 2015). The seeds in these treatments were rinsed with water and planted in nursery bags as shown in plate 3. The nursery bags were labeled according to the type of seeds, treatment used and time variation. The planted seeds were observed for a period of 30 days and records were taken daily.

**Plate 1 & 2:**
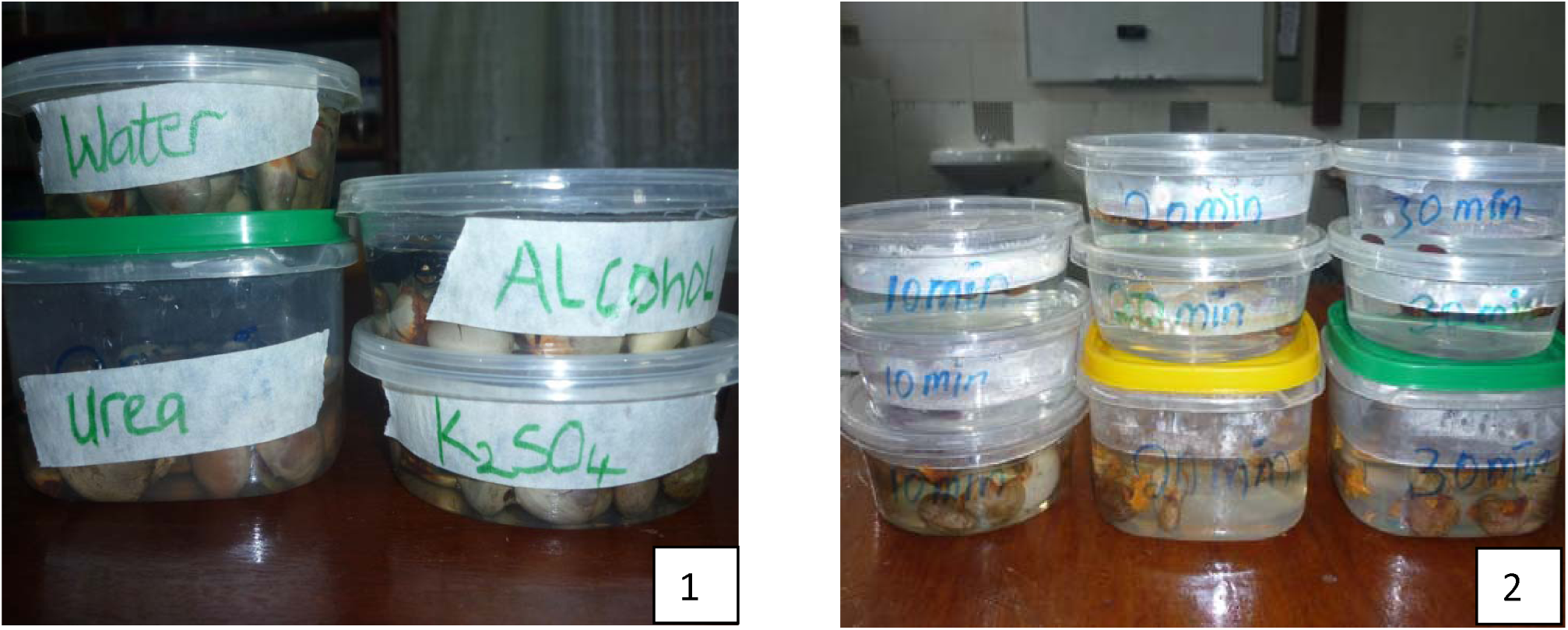
Seeds soaked in different treatments of water Urea, alcohol and potassium sulphate

**Plate 3:**
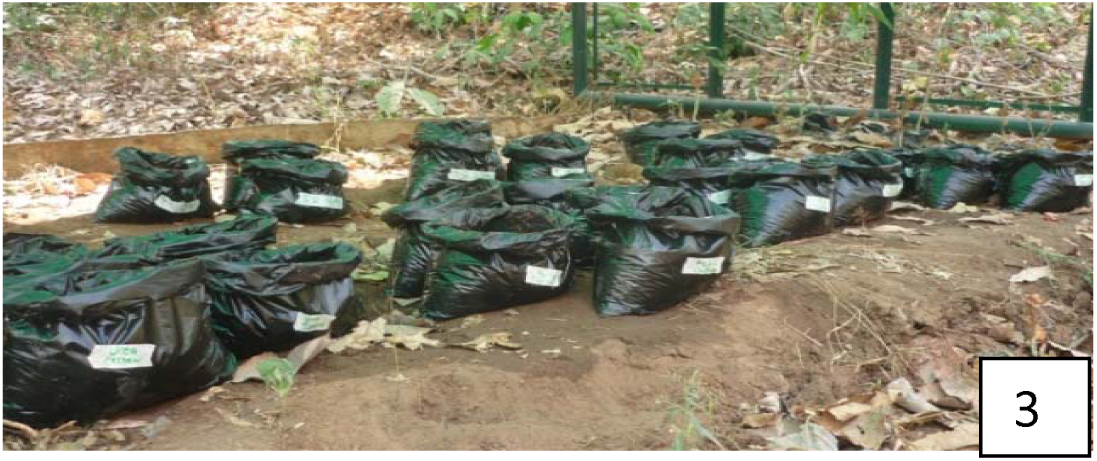
Seeds planted in polythene bags

## RESULT

White arrows indicate seeds of *Jatropha curcas, Tamarindus indica, Anona muricata and Anacardium occidentale* treated with Nitric acid having seedlings with luxuriant growth after 30 days, *Artocarpus heterophyllus* did not survive in it (plate 4 and table 1).

**Plate 4:**
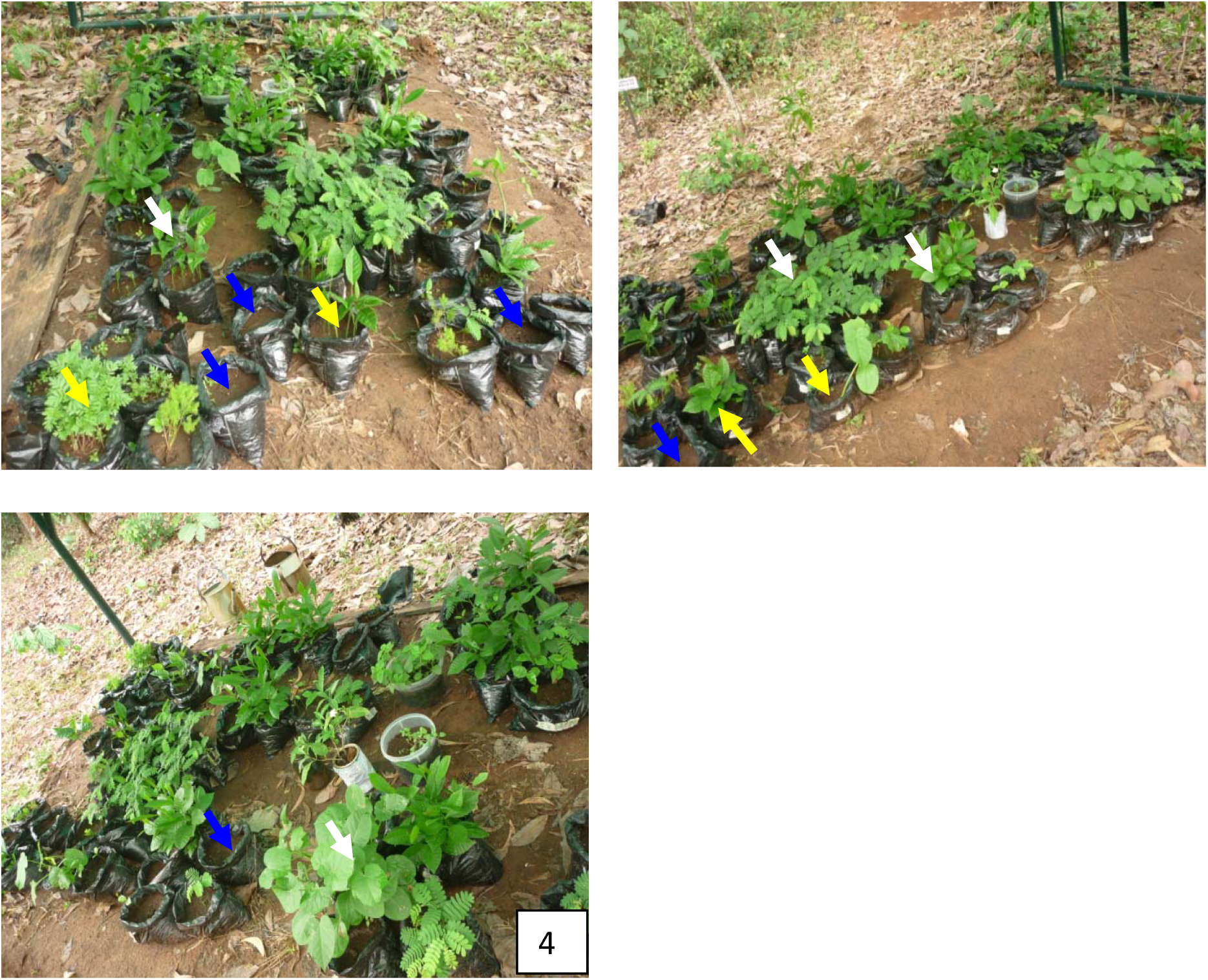
Seedlings after thirty days (30) of planting

**Table 1:** Result Effect of the treatments on seeds used

| Seeds | 65% HNO<sub>3</sub><br>10 mins | 65% HNO<sub>3</sub><br>20 mins | 65% HNO<sub>3</sub><br>30 mins | 65% H<sub>2</sub>SO<sub>4</sub><br>10 mins | 65% H<sub>2</sub>SO<sub>4</sub><br>20 mins | 65% H<sub>2</sub>SO<sub>4</sub><br>30 mins | H<sub>2</sub>O 24 hrs | 0.5% K<sub>2</sub>SO<sub>4</sub><br>24 hrs | 0.5% CH<sub>4</sub>N<sub>2</sub>O<br>24 hrs | 43% C<sub>2</sub>H<sub>6</sub>O<br>24 hrs | Control |
| --- | --- | --- | --- | --- | --- | --- | --- | --- | --- | --- | --- |
| ***Anacardium occidentale*** |  |  |  |  |  |  |  |  |  |  |  |
| No of seeds used | 10 | 10 | 10 | 10 | 10 | 10 | 10 | 10 | 10 | 10 | 10 |
| No of germinated seeds | 10 | 5 | 7 | 8 | 7 | 7 | 1 | 5 | 9 | 2 | 8 |
| Days taken to germinate | 15 | 17 | 13 | 16 | 16 | 15 | 19 | 15 | 16 | 17 | 19 |
| ***Jatropha curcas*** |  |  |  |  |  |  |  |  |  |  |  |
| No of seeds used | 10 | 10 | 10 | 10 | 10 | 10 | 10 | 10 | 10 | 10 | 10 |
| No of germinated seeds | 8 | 0 | 0 | 5 | 3 | 0 | 8 | 10 | 0 | 0 | 9 |
| Days taken to germinate | 6 | 0 | 0 | 7 | 6 | 0 | 9 | 9 | 0 | 0 | 6 |
| ***Tamarindus indica*** |  |  |  |  |  |  |  |  |  |  |  |
| No of seeds used | 10 | 10 | 10 | 10 | 10 | 10 | 10 | 10 | 10 | 10 | 10 |
| No of germinated seeds | 10 | 9 | 10 | 8 | 4 | 0 | 10 | 10 | 9 | 1 | 10 |
| Days taken to germinate | 12 | 17 | 14 | 19 | 19 | 0 | 6 | 11 | 14 | 17 | 16 |
| ***Annona muricata*** |  |  |  |  |  |  |  |  |  |  |  |
| No of seeds used | 10 | 10 | 10 | 10 | 10 | 10 | 10 | 10 | 10 | 10 | 10 |
| No of germinated seeds | 8 | 9 | 7 | 5 | 8 | 7 | 6 | 6 | 2 | 0 | 7 |
| Days taken to germinate | 19 | 22 | 23 | 20 | 22 | 22 | 19 | 23 | 24 | 0 | 24 |
| ***Artocarpus heterophyllus*** |  |  |  |  |  |  |  |  |  |  |  |
| No of seeds used | 10 | 10 | 10 | 10 | 10 | 10 | 10 | 10 | 10 | 10 | 10 |
| No of germinated seeds | 0 | 0 | 0 | 0 | 0 | 0 | 7 | 9 | 2 | 0 | 9 |
| Days taken to germinate | 0 | 0 | 0 | 0 | 0 | 0 | 14 | 15 | 24 | 0 | 18 |

Yellow arrows indicate seeds of *Jatropha curcas, Artocarpus heterophyllus, Annona muricata and Tamarindus indica* treated with water having seedling with variations in growth and survival after 30 days. *Anacardium occidentale* did not survive well in this treatment, because most of the seeds started fermenting after 24 hours thus it produced only one seedling after planting the seeds (plate 4 and table 1).

Blue arrow indicates seeds of *Anacardium occidentale, Jatropha curcas, Artocarpus heterophyllus, Annona muricata and Tamarindus indica* treated with Nitric acid, Sulphuric acid, Urea and Ethanol. *Anacardium occidentale* survived in nitric acid, sulphuric acid and urea but it produced only 2 seedlings in ethanol treatment. *Jatropha curcas* only survive in nitric and sulphuric acid at limited time of 10–20 minutes treatment but did not survive in urea and ethanol. *Artocarpus heterophyllus* did not survive in nitric acid, sulphuric acid and ethanol due to its tin seed coat that could be rapidly degraded by acids and alcohol but 2 seedlings was produced in the treatment with urea. *Annona muricata* and *Tamarindus indica* survived in nitric acid, sulphuric acid and urea. *Tamarindus indica* produced one seedling in ethanol treatment but *Annona muricata* did not survived in it (plate 4 and table 1).

The whole chemical treatments process for *Anacardium occidentales* produced viable seedlings, although water produced only one viable seedling and two for ethanol (43% C_2_H_6_O for 24 hours), which produced two viable seedlings (as seen in Table 1) with germination period of 19 and 17 respectively. The set germination period for the controlled experiment was 19 days. Water has same germination period with the set value of the controlled experiment. The reason for decrease in number of seedling survival in water is due to the formation of foam (lather) which might be as a result of fermentation causing decay of the embryo. The best treatment for *Anacardium occidentales* is Nitric acid (65% HNO_3_ for 10 minutes) which produced high numbers of seedlings or viable seeds (10) and a germination period of 15 days as supported by the work of Yücel (2000). Oyewole & Koffa, (2010) work on the response of sprouting in cashew nut due to storage length, nut size, nut soaking and their interactions. They discovered that nuts presoaked in water for 24 or 36 hours before sowing was observed to give the best result for soaking treatment and therefore recommend it to farmers involved in large scale cashew farming. This result does not fit into our experiments as fermentation was noticed after 24 hours of soaking cashew seed.

**Plate 5:**
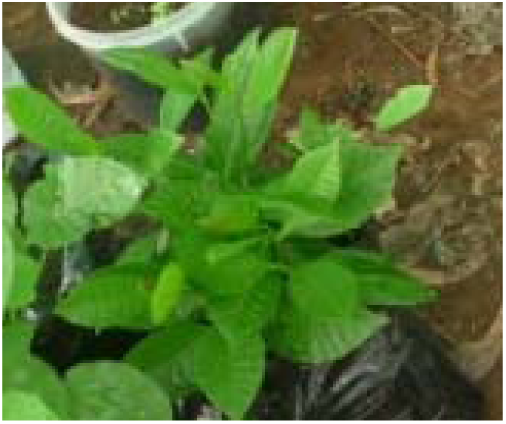
Seedlings of *Anacardium occidentale*

**Plate 6:**
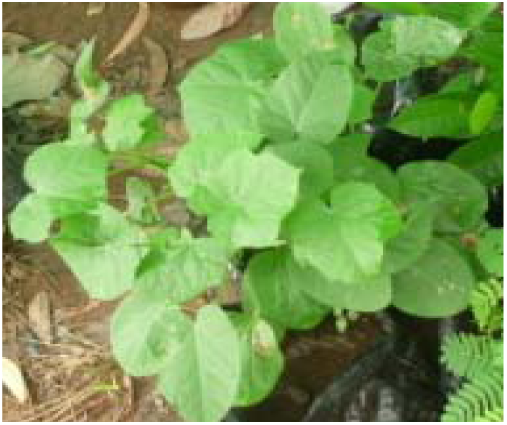
Seedlings of *Jatropha curcas*

**Plate 7:**
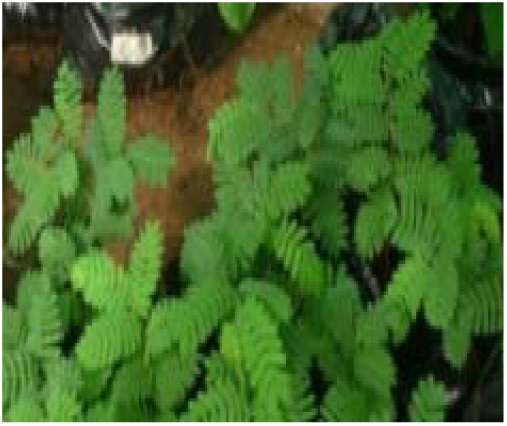
Seedlings of *Tamarindus indica*

**Plate 8:**
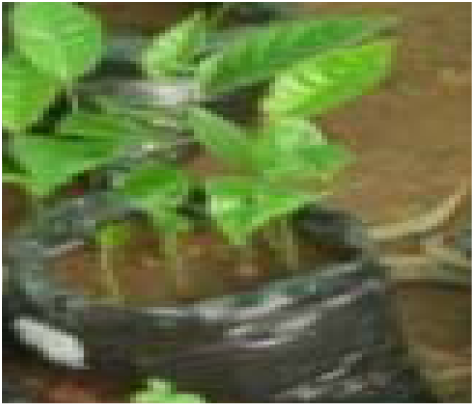
Seedlings of *Annona muricata*

**Plate 9:**
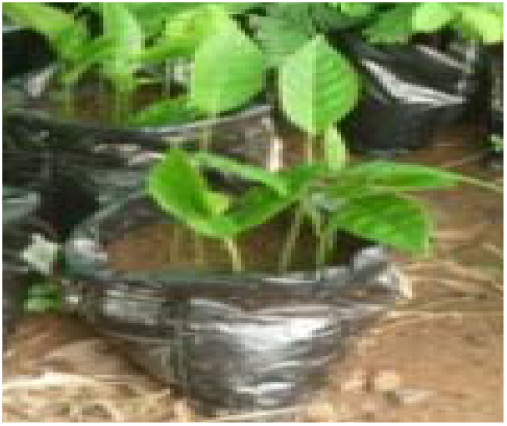
Seedlings of *Artocarpus heterophyllus*

The seeds of *Jatropha curcas* did not survive in Urea and ethanol (0.5% CH_4_N_2_O and 43% C_2_H_6_O soaked for 24 hours). Though they survive the Nitric and sulphuric acid treatments at 10 minutes but not above it as supported by the work done by Davis *et al.* (1991) for the improvement of seedling emergence of *Lupinus texensis* through acid scarification. Seed survival decreases as the time increases for the acid treatments (20–30 minutes) as seen in Table 1. Many of the survived acid treated seeds had delay germination as against the control. There was no significant difference between acid treated seeds (Nitric acid) and the control seeds germination time. The luxuriant growth of *Jathropha curcas* might be due to the presence of microorganism as reported in the work of Jha & Saraf, (2011), on the effect of plant growth promoting rhizobacteria on seed germination, behaviour and seedling vigor of *Jatropha curcas* seed inoculated with five strains of bacteria isolates; *Brevibacillus brevis* MS1, *Enterobacter aerogenes* MS2, *Bacillus licheniformis* MS3, *Micrococcus* sp. MS4 and *Acienetobacter calcoaceticus* MS5. The highest germination percentage was observed in MS5 (63.33 %) and the germination capacity (80 %) followed by MS3 (60 %) with germination capacity (73.33 %) while maximum seedling vigor index (929.05) was reported in MS5 followed by MS3 (895.8). The seedling vigor reported in our work in treatment with Nitric acid for 10 minute may be due to the presence of residual nitrogen compound from the treated seed couple with the activities of soil microorganism which naturally fix nitrogen compound to plant root in which the five strains of microorganism mentioned in the work of Jha & Saraf, (2011) above falls into that category.

The whole chemical treatments for *Tamarindus indica* was favourable except ethanol (43% C_2_H_6_O for 24 hours) which produce only one seedling and took one extra day (17 day) to germinate above the set control value (16 days) and also sulphuric acid (65% H_2_SO_4_ for 30 minutes) which does not produce any seedling. Nitric acid (65% HNO_3_ for 10 minutes) and potassium sulphate (0.5% K_2_SO_4_ for 24 hours) produce seedlings at periods with close range (12 and 11 days) but the best treatment for *Tamarindus indica* is water which recorded no seed mortality with six days of germination which is better than the control value (16) as seen in Table 1. Information available on breaking seed coat dormancy in *Tamarindus indica is* very meager. Naikawadi, (2016) soaked seed of *Tephrosia purpurea* with 75% H_2_SO_4_ for 2.5 minutes, 75% HCl for 5 minutes, Indole-3-acetic (IAA) and Gibbrellic acid (GA3), 5.0 mM GA3 for 24 hours and discovered that seeds soaked in IAA and GA3, 5.0 mM GA3 for 24 hours treatment gives better germination and other treatments produced fairly good seedlings. Though *Tephrosia purpurea* and *Tamarindus indica* are not closely related in terms of their generic name but they are closely related at their family level. This may suggest the reason why their seeds have similar responses to acids and water.

Similar work done by Tigabu & Odén, (2001) on the Effect of scarification, gibberellic acid and temperature on seed germination of *Albizia grandibracteata* and *Albizia gummifera* from Ethiopia, which belongs to the same family and subfamily with *Tamarindus*. They soaked these two species of *Albizia* in sulpuric acid, gibberelic acid and hot water. The whole treatment improves the germination and vigour of the plant species which shares specific pod and seed characteristics with *Tamarindus indica.*

The seeds of *Annona muricata* did not survive in ethanol (0.5% CH_4_N_2_O and 43%) and only 2 seeds survived in urea (0.5% CH_4_N_2_O) but other treatments produced viable seedlings. Nitric acid (65% HNO_3_ for 10 minutes) and water has favourable germination period of 19 days against the set period of the control experiment which is 24 days (See Table 1). Similar result was obtained in work done by Chagas *et al.*, (2013) which was geared at breaking the dormancy in sugar apple (*Annona muricata*) seeds using physical and chemical method. They found out that scarification and soaking in water for 24 h is economical and has a lot of practical applications which concur with the 19 days germination period in our experiment.

The seeds of *Artocarpus heterophyllus* only survived in Urea, potassium sulphate and water (0.5% CH_4_N_2_O, 0.5% K_2_SO_4_ and H_2_O). It does not survive in high concentration of the acids used in this experiment due to its thin seed coat. Water and potassium sulphate are the best treatments for *Artocarpus heterophyllus* as they produce viable seedlings with short germination period of 14 and 15 days against the 18 days of the control experiment as seen in Table 1. The results of this finding are in conformity with the findings of LAL, (2015), in the area of reduction of germination days by soaking seeds of *Artocarpus heterophyllus* in growth regulators and thiourea.

## CONCLUSION

The best treatment for *Anacardium occidentale* is Nitric acid which produced high numbers of seedlings or viable seeds and low germination period. It is better to plant *Jathropha curcas* in the cultural way of planting it than using chemical treatments. Nitric acid and water are the best treatments for *Annona muricata* and *Tamarindus indica.* Water and potassium sulphate are the best treatments for *Artocarpus heterophyllus,* Seeds treated with Nitric acid produced healthy seedlings after 30 days than the rest treatments.

## REFERENCES

Baskin CC. 2003. Breaking physical dormancy in seeds – Focussing on the lens. New Phytologist 158: 229–232.

Baskin CC, Baskin JM. 1998. Seeds: ecology, biogeography, and evolution of dormancy and germination. Seeds: ecology, biogeography, and evolution of dormancy and germination.: xiv + 666 pp.

Baskin JM, Baskin CC. 2004. A classification system for seed dormancy. Seed Science Research 14.

Bentsink L, Koornneef M. 2008. Seed Dormancy and Germination. The Arabidopsis Book 6: e0119.

Bewley JD, Black M. 1994. Seeds: physiology of development and germination. Plenum Press.

Casal JJ, Sánchez RA. 1998. Phytochromes and seed germination. Seed Science Research 8.

Chagas PC, Sobral STM, Oliveira RR de, Chagas EA, Pio R, Santos VA dos. 2013. Physical and chemical methods to breach seed dormancy of sugar apple. Amazonian Journal of Agricultural and Environmental Sciences 56: 1–6.

Davis TD, George SW, Upadhyaya A, Parsons J. 1991. Improvement of Seedling Emergence of Lupinus texensis Hook. Following Seed Scarification Treatments. Journal of Environmental Horticulture 9: 17–21.

El-Siddig K, Ebert G, Lüdders P. 2001. A comparison of pretreatment methods for scarification and germination of Tamarindus indica L. seeds. Seed Science and Technology 29: 271–274.

Fenner M, Thompson K. 2005. The ecology of seeds.

Finch-Savage WE, Clay HA. 1997. The influence of embryo restraint during dormancy loss and germination of Fraxinus excelsior seeds. In: Basic and Applied Aspects of Seed Biology. 245–253.

Finch-Savage WE, Leubner-Metzger G. 2006. Seed dormancy and the control of germination. New Phytologist 171: 501–523.

Hilhorst HWM. 1995. A critical update on seed dormancy. I. Primary dormancy. Seed Science Research 5: 61–73.

Jacobsen J V., Pressman E. 1979. A structural study of germination in celery (Apium graveolens L.) seed with emphasis on endosperm breakdown. Planta 144: 241–248.

Jha KS, Saraf M. 2011. Effect of Plant Growth Promoting Rhizobacteria on Seed Germination Behaviour and Seedling Vigor of Jatropha curcas. International Journal of Biotechnology & Biosciences 1: 161–169.

Kucera B, Cohn MA, Leubner-Metzger G. 2005. Plant hormone interactions during seed dormancy release and germination. Seed Science Research 15: 281–307.

Lal I. 2015. “Response of Jackfruit (Artocarpus heterophyllus Lam.) Seeds to Plant Growth Regulators and Thiourea”.

Li B, Foley ME. 1997. Genetic and molecular control of seed dormancy. Trends in Plant Science 2: 384–389.

Naikawadi VB. 2016. SEED GERMINATION OF IMPORTANT MEDICINAL PLANT TEPROSIA PURPUREA (LINN), PERS. International Research Journal of Natural and Applied Sciences 3.

Nikolaeva MG. 1969. Physiology of deep dormancy in seeds. Leningrad, Russia, Izdatel’stvo ‘Nauka’. (Translated from Russian by Z. Shapiro, National Science Foundation, Washington, DC.): 219.

Nikolaeva MG. 2004. On criteria to use in studies of seed evolution. Seed Science Research 14: 315–320.

Offord CA, Meagher PF, Australian Network for Plant Conservation., Australian Seed Conservation and Research. 2009. Plant germplasm conservation in Australia: strategies and guidelines for developing, managing and utilising ex situ collections. Australian Network for Plant Conservation (ANPC), in partnership with Australian Seed Conservation and Research (AuSCaR).

Oyewole CI, Koffa KJ. 2010. Effect of Storage, Size of Nut and Soaking Length on Sprout Emergence in Cashew. www.thaiagj.org Thai Journal of Agricultural Science 43: 39–45.

Purohit S, Nandi SK, Palni LMS, Giri L, Bhatt A. 2015. Effect of Sulfuric Acid Treatment on Breaking of Seed Dormancy and Subsequent Seedling Establishment in Zanthoxylum armatum DC: An Endangered Medicinal Plant of the Himalayan Region. National Academy Science Letters 38: 301–304.

Rooser P. 2017. Seed science and technology. Larsen and Keller Education.

Sanchez R, Mella R. 2004. The exit from dormancy and the induction of germination: physiological and molecular aspects. In: Benech-Arnold, Roberto L.; Sanchez RA, ed. Handbook of seed physiologyl: applications to agriculture. Binghamton, New York: Food Products Press, 221–243.

Thompson K. 2000. The Functional Ecology of Soil Seed Banks. Seeds: The Ecology of Regeneration in Plant Communities: 215–236.

Tigabu M, Odén PC. 2001. Effect of scarification, gibberellic acid and temperature on seed germination of two multipurpose Albizia species from Ethiopia. Seed Science and Technology 29: 11–20.

Vleeshouwers LM, Bouwmeester HJ, Karssen CM. 1995. Redefining Seed Dormancy: An Attempt to Integrate Physiology and Ecology. The Journal of Ecology 83: 1031.

Yücel E. 2000. Effects of different salt (NaCl), nitrate (KNO3) and acid (H2SO4) concentrations on the germination of some Salvia species seeds. Seed Science and Technology 28: 853–860.

